# An insight to HTLV-1-associated myelopathy/tropical spastic paraparesis (HAM/TSP) pathogenesis; evidence from high-throughput data integration and meta-analysis

**DOI:** 10.1101/754697

**Authors:** Sayed-Hamidreza Mozhgani, Mehran Piran, Mohadeseh Zarei-Ghobadi, Mohieddin Jafari, Talat Mokhtari-Azad, Seyed-Mohammad Jazayeri, Majid Teymoori-Rad, Narges Valizadeh, Hamid Farajifard, Mehdi Mirzaie, Azam Khamseh, Houshang Rafatpanah, Seyed-Abdolrahim Rezaee, Mehdi Norouzi

**Author notes:** S-H.M, M. Z-G, M.P, and M.J are co-first authors of the paper. **Corresponding Authors:** Seyed-Abdolrahim Rezaee, Mehdi Norouzi.

## Abstract

Human T-lymphotropic virus 1-associated myelopathy/tropical spastic paraparesis (HAM/TSP) is a progressive disease of the central nervous system that affected significantly spinal cord, nevertheless, the pathogenesis pathway. This study aimed to employ high throughput meta-analysis to find major genes involved in the pathogenesis of HAM/TSP. High-throughput statistical analyses identified 385, 49, and 22 differentially expressed genes for normal *vs.* ACs, normal *vs.* HAM/TSP and ACs *vs.* HAM/TSP groups, respectively. STRING and further network analyses highlighted 32, 29, and 13 hub genes for normal *vs.* ACs, normal *vs.* HAM/TSP, and ACs *vs.* HAM/TSP groups, respectively. Biological network analyses indicated the involvement of hub genes in the HAM/TSP group in many vital pathways like apoptosis and immune pathways. Moreover, the meta-analysis results disclosed three major genes including STAT1, TAP1, and PSMB8 which have function role in HAM/TSP progression. Real-time PCR revealed the meaningful down-regulation of STAT1 in HAM/TSP samples than AC and normal samples (*P*=0.01 and *P*=0.02, respectively), up-regulation of PSMB8 in HAM/TSP samples than AC and normal samples (*P*=0.04 and *P*=0.01, respectively), and down-regulation of TAP1 in HAM/TSP samples than those in AC and normal samples (*P*=0.008 and *P*=0.02, respectively). No significant difference was found among three groups in terms of percentage of T helper and cytotoxic T lymphocytes (*P*= 0.55 and *P*=0.12). Our results confirm that STAT1, TAP1, and PSMB8 are three important genes which their expressions levels were changed in three different groups. These proteins in association with other proteins can involve in the immune and apoptosis pathways.

## Introduction

HTLV-associated myelopathy/tropical spastic paraparesis (HAM/TSP) is a chronic neurodegenerative disease with progressive characteristics that disturbs the functioning of the sensory and motor nerves (Andrade et al., 2013). Indeed, infection with HTLV-1 can lead to asymptomatic carrier (AC) state or two diseases including Adult T-Cell Leukemia Lymphoma (ATLL) or/and HAM/TSP (Mozhgani et al., 2018b).

About 10-20 million people worldwide have been infected with HTLV-1 (Trevino et al., 2012). Endemic areas include the Middle East, Japan, the Caribbean basin, Central Africa, the Melanesian Islands, and South America. Only 2-5 % of those infected with the virus develop HAM/TSP (Ahmadi Ghezeldasht et al., 2013;Shoeibi et al., 2013).

Patients with HAM/TSP often have symptoms such as back pain, stiffness, and pain in the lower limbs, urinary frequency, and progressive weakness. Mild cognitive impairment is also common. The clinical signs of the disease imitate multiple sclerosis when the spinal cord is involved, such that sick people need walking aids after one year of illness (Nakagawa et al., 1995).

HTLV-1 may weaken or impair the immune system, which results in autoimmunity to neurons. It also provides an immunosuppressive microenvironment that authorizes the HTLV-1 infected cells to escape host immune response and causes HTLV-1-associated diseases (Kchour et al., 2013).

Studies on HTLV-1 as a factor that deregulates the host’s immune system has lasted for many years and has sometimes yielded polemical results. After years of study on HAM/TSP treatment, it is still a challenge for clinicians (Hermine et al., 1998). Therefore, identifying prognostic biomarkers that implicated in the pathogenesis is vital to understand the development and progression of a disease, as well as its diagnosis and treatment. Since now, different genes that are involved in mTOR, NF-kappa B, PI3K, and MAPK signaling pathways have been known in the HAM/TSP cases. Also, apoptosis can occur in the cell nucleus of the HAM/TSP patients (Moriuchi et al., 1999;Ottoson et al., 2001;Mozhgani et al., 2018b).

Microarray technology can simultaneously measure tens of thousands of genes from different tissue samples in a high-throughput and cost-effective manner (Schena et al., 1995). However, the results may be irreproducible (Ntzani and Ioannidis, 2003) or be influenced by the data perturbations (Ein-Dor et al., 2005;Michiels et al., 2005). One possible solution to find robust information is the integration of multiple data which is called meta-analysis (Gieger et al., 2008;Zhang et al., 2013;Li et al., 2014;Pena et al., 2014). To this end, various statistical procedures are employed to combine and analyze the results of the independent studies. Meta-analysis increases the validity of the results and makes the possible estimation of gene expression differences (Ramasamy et al., 2008b).

In this study, we integrated datasets to find gene signatures by network analysis of differentially expressed genes. The results specified the genes and pathways, which possibly have critical roles in the development of HAM/TSP pathogenesis. Flow cytometry was employed to determine the ratio of CD4+ to CD8+ and better understanding pathogenesis of the virus. Moreover, the real-time PCR confirmed different expressions of the determined genes in the HAM/TSP cases versus AC and normal subjects.

## Materials and Methods

### Database searching and identification of eligible studies

We searched the Gene Expression Omnibus (http://www.ncbi.nlm.nih.gov/geo/) and ArrayExpress (https://www.ebi.ac.uk/arrayexpress/) by end of 2018 to find datasets reporting the expression levels of miRNA and mRNA in the HAM/TSP and AC subjects. The keywords including Human T-lymphotropic virus 1-associated myelopathy/tropical spastic paraparesis, HTLV-1, TSP, HAM/TSP, asymptomatic carrier, AC, ACs were used to find the relevant reports. The inclusion criteria were homo sapiens datasets in which peripheral blood mononuclear cells (PBMCs) were analyzed in AC or HAM/TSP groups. The normal samples were also considered in these groups. The exclusion criteria were studies which performed on Mus musculus, cell line, and samples other than PMBCs. Two independent investigators searched and gathered data from each included study. The quality and consistency of the studies were evaluated by the R package MetaQC (Kang et al., 2012). Finally, the obtained data were classified into three groups named as ACs *vs.* normal, HAM/TSP *vs.* normal, and HAM/TSP *vs.* ACs.

### Pre-processing and meta-analysis

The expression data in each group were background corrected, logarithmically transformed (base 2) and quantile normalized using the Affy package implemented in R programming language (www.r-project.org). The datasets were integrated individually at miRNA and mRNA levels using random effect method (REM) and then differentially expressed miRNAs (DEMs) and differentially expressed genes (DEGs) were identified, respectively. False discovery rate (FDR) of <0.05 was considered a significant difference. The target genes of each DEMs were obtained using miRDB (http://mirdb.org) and then integrated super-horizontally with DEGs. The common genes were considered for further analysis.

### Super- and sub-networks construction

To construct the super-network comprises protein-protein interactions (PPIs) in each group, the STRING database version 10.5 was employed (Szklarczyk et al., 2016). PPIs networks were analyzed in terms of degree and betweenness centrality by NetworkAnalyzer in Cytoscape 3.5.1. The genes with higher aforementioned criteria were considered as hub genes. Afterward, the hub genes were utilized to construct the sub-networks. The sub-networks were visualized by Gephi (0.9.1) (Bastian et al., 2009).

### Module finding and pathways analysis

The sub-network clustering was implemented using the fast unfolding clustering algorithm in Gephi (Mozhgani et al., 2018b). The biologically meaningful modules were then chosen. To find the KEGG pathways in which hub genes are involved, Enrichr web tool was employed (Kuleshov et al., 2016). Ten top pathway terms that had higher combined scores were selected for further interpretations.

### Patient population and Sample collection

Blood samples were collected from eight patients with ACs, eight patients with HAM/TSP, and eight normal samples who referred to the neurology department of Ghaem Hospital, Mashhad University of Medical Sciences (MUMS). All specimens were collected after acquiring informed consent from the patient’s guardians. Two trained neurologists affirmed the diagnosis of HAM/TSP according to WHO criteria. All contributors had seropositive test for HTLV-1 by enzyme-linked immunosorbent assay (ELISA, Diapro, Italy). The results of serology were confirmed by PCR (Rafatpanah et al., 2016). The participants had no history of treatment with IFNs. This study was approved by the Ethics Committee of Biomedical Research at MUMS (IR.TUMS.SPH.REC.1396.242).

### Flow cytometry analysis

To determine T helper and cytotoxic cells populations in HAM/TSP, ACs and normal groups; PerCP anti CD3 antibody (bio legend company cat no: 344813), Phicoerythrin (PE) anti CD4 antibody (bio legend company cat no: 317409) and PE anti CD8 antibody (bio legend company cat no: 301007) were used. Fresh peripheral blood samples were treated by lysis buffer for destroying the red blood cells and platelets. Samples were analyzed on a FACS caliber Becton Dichinson. All analyses were done in the lymphocyte gate.

### HTLV-1 Proviral Load

Peripheral blood mononuclear cells (PBMCs) were isolated from EDTA-treated blood samples using Ficoll density gradient medium (Cedarlane, Hornsby, ON, Canada). The commercial blood mini kit (Qiagen, Germany) was applied to extract DNA from PBMCs. In order to measure the PVL of HTLV-I in PBMCs, a real-time PCR using a commercial real-time-based absolute quantification kit (HTLV-1 RG; Novin Gene, Karaj, Iran) was performed (Boostani et al., 2015).

### Quantitative real-time PCR

Total RNA was extracted from fresh PBMCs using TriPure isolation reagent (Roche, Germany) according to the manufacturer’s instructions. Double-stranded cDNA was synthesized using the RevertAid TM first strand cDNA synthesis kit (Fermentas, Germany). Following primers and probes were designed and used to determine the expression levels of STAT1, PSMB8, TAP1 : STAT1 (forward primer: 5□-AACATGGAGGAGTCCACCAATG-3□, reverse primer: 5□-GATCACCACAACGGGCAGAG-3□ and TaqMan probe: FAM-TCTGGCGGCTGAATTTCGGCACCT -BHQ1), PSMB8 (forward primer: 5□-GTTCAGATTGAGATGGCCCATG-3□, reverse primer: 5□-CGTTCTCCATTTCGCAGATAGTAC-3□ and TaqMan probe: FAM-CCACCACGCTCGCCTTCAAGTTCC -BHQ1), TAP1 (forward primer: 5□-TACCGCCTTCGTTGTCAGTTATG-3□, reverse primer: 5□-GAGCCCAGGCAGCCTAGAAG-3□ and TaqMan probe: Fam-CGCACAGGGTTTCCAGAGCCGCC-BHQ1). The primers and probes of Tax and HBZ were synthesized according to published data (Mozhgani et al., 2018a). The relative 2 standard curves real-time PCR was carried out on the cDNA samples using TaqMan master mix (Takara, Otsu, Japan) and a Q-6000 machine (Qiagen, Germany). The GAPDH gene was employed as a housekeeping gene to normalize mRNA expression levels and also to control the error between samples (Boostani et al., 2015;Jafarian et al., 2017).

### Statistical analysis

Statistical analysis was carried out using GraphPad Prism Software Version 7 (GraphPad software, Inc). Quantitative data were expressed as mean ± SEM and percentages. The comparisons between various groups were accomplished using ANOVA. Pearson’s or Spearman’s tests were used for the analysis of the correlation between variables. The outcomes were considered significant if *P* ≤ 0.05.

## Results

### Studies included in the meta-analysis

According to our criteria, 16 studies were found in the GEO repository datasets which were performed at mRNA or miRNA levels. After quality control done by MetaQC packages, seven (GSE29312, GSE29332, GSE46518, GSE52244, GSE55851, GSE11577, GSE46345), three (GSE19080, GSE29312, GSE29332), and four (GSE38537, GSE29312, GSE29332, GSE19080) mRNA and miRNA datasets were of high quality for further analyses of normal *vs.* ACs, normal *vs.* HAM/TSP, and ACs *vs.* HAM/TSP groups, respectively (Table 1).

**Table 1.** Selected studies included in the meta-analysis.

| Eligible studies in the meta-analysis |  |  |
| --- | --- | --- |
| Row | Normal *vs.* ACs | Type |
| 1 | GSE29312 | Expression profiling by array |
| 2 | GSE29332 | Expression profiling by array |
| 3 | GSE46518 | Expression profiling by array |
| 4 | GSE52244 | Expression profiling by array |
| 5 | GSE55851 | Expression profiling by array |
| 6 | GSE11577 | Non-coding RNA profiling by array |
| 7 | GSE46345 | Non-coding RNA profiling by array |
| Row | Normal *vs.* TSP | Type |
| 1 | GSE19080 | Expression profiling by array |
| 2 | GSE29312 | Expression profiling by array |
| 3 | GSE29332 | Expression profiling by array |
| Row | ACs *vs.* TSP | Type |
| 1 | GSE38537 | Expression profiling by array |
| 2 | GSE29312 | Expression profiling by array |
| 3 | GSE29332 | Expression profiling by array |
| 4 | GSE19080 | Expression profiling by array |

### Differentially Expressed genes and miRNAs

A total of four miRNAs including hsa-mir-218, hsa-mir-206, hsa-mir-31, and hsa-mir-34A were identified as DEMs between normal and ACs groups. The target genes of the mentioned DEMs were further identified in miRDB. Among 3001 genes, 207 genes with rank_product<20 were selected and added to 180 DEGs obtained across microarray datasets. After removing duplicate genes, 385 DEGs were specified for super-network analysis. Also, a total of 49 and 22 genes were identified to be DEGs for normal *vs.* HAM/TSP and ACs *vs.* HAM/TSP groups, respectively.

### Protein-protein interactions networks (PPINs)

To explore more information about the relationships between the DEGs, PPINs were constructed by STRING. Firstly, all DEGs were considered for building super-PPINs in each group. The networks were analyzed in terms of topology and centrality parameters. The nodes with higher degree and betweenness were selected as hub genes. From these analyses, 32, 29, and 13 hub genes were specified for normal *vs.* ACs, normal *vs.* HAM/TSP, and ACs *vs.* HAM/TSP groups, respectively (Table 2). Afterward, the hub genes were again submitted to STRING to find sub-networks as shown in Figure 1. The networks consist of 32 nodes and 106 edges for normal *vs.* ACs (Figure 1a), 29 nodes and 96 edges for normal *vs.* TSP (Figure 1b), 13 nodes and 18 edges for ACs *vs.* TSP (Figure 1c). The size and color of each node is representative of the degree level. The red color and bigger size indicate the higher degree of the node, which in turn shows the important role of that node in the network. The top ten genes according to degree and betweenness centrality analysis in the studied groups, as well as common genes found between these two criteria in each group are shown in Table 3.

**Table 2.**
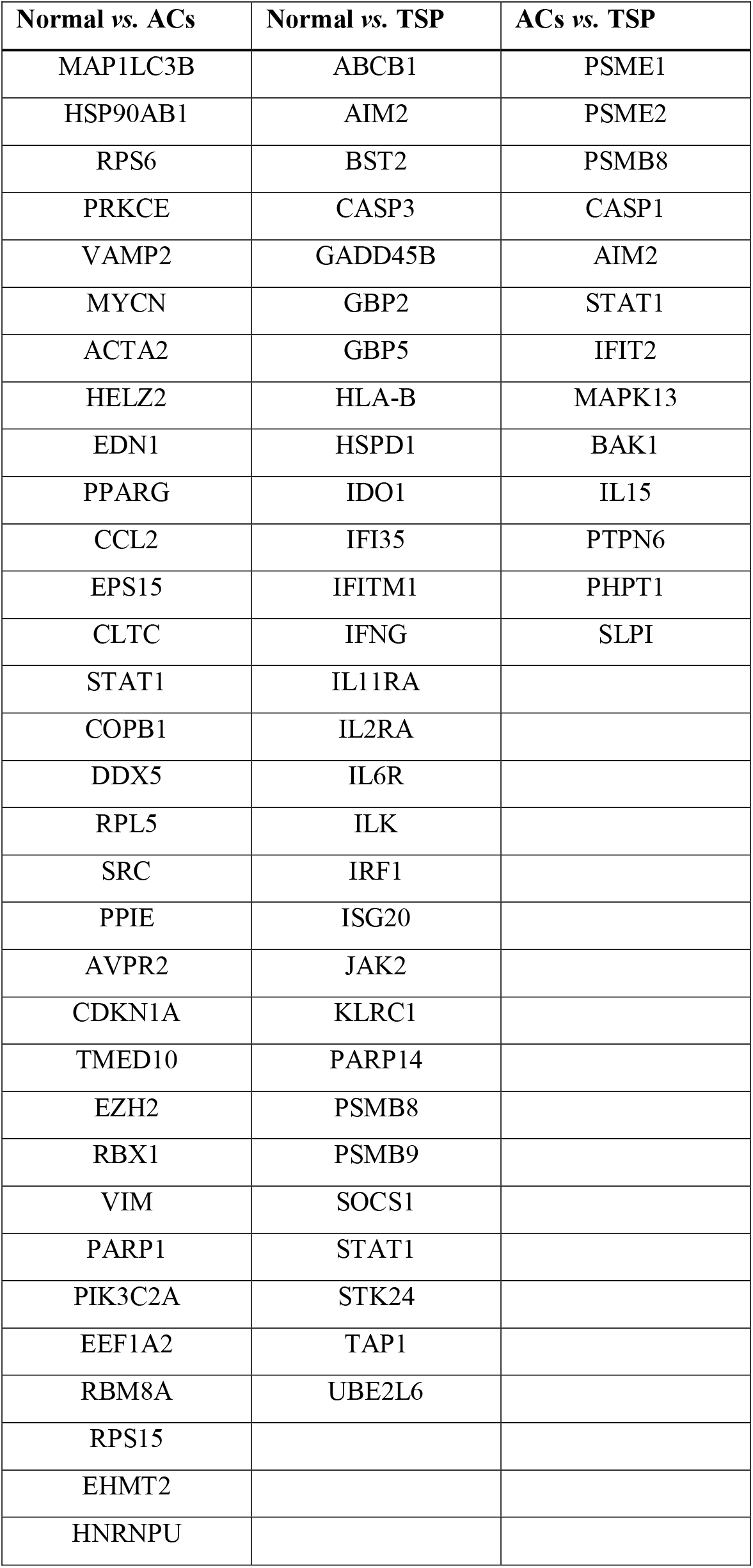
Hub differentially expressed genes including higher degree and betweenness in the sub networks of Normal *vs.* ACs, Normal *vs.* HAM/TSP, and ACs *vs.* HAM/TSP groups.

| Normal vs. ACs | Normal vs. TSP | ACs vs. TSP |
| --- | --- | --- |
| MAP1LC3B | ABCB1 | PSME1 |
| HSP90AB1 | AIM2 | PSME2 |
| RPS6 | BST2 | PSMB8 |
| PRKCE | CASP3 | CASP1 |
| VAMP2 | GADD45B | AIM2 |
| MYCN | GBP2 | STAT1 |
| ACTA2 | GBP5 | IFIT2 |
| HELZ2 | HLA-B | MAPK13 |
| EDN1 | HSPD1 | BAK1 |
| PPARG | IDO1 | IL15 |
| CCL2 | IFI35 | PTPN6 |
| EPS15 | IFITM1 | PHPT1 |
| CLTC | IFNG | SLPI |
| STAT1 | IL11RA |  |
| COPB1 | IL2RA |  |
| DDX5 | IL6R |  |
| RPL5 | ILK |  |
| SRC | IRF1 |  |
| PPIE | ISG20 |  |
| AVPR2 | JAK2 |  |
| CDKN1A | KLRC1 |  |
| TMED10 | PARP14 |  |
| EZH2 | PSMB8 |  |
| RBX1 | PSMB9 |  |
| VIM | SOCS1 |  |
| PARP1 | STAT1 |  |
| PIK3C2A | STK24 |  |
| EEF1A2 | TAP1 |  |
| RBM8A | UBE2L6 |  |
| RPS15 |  |  |
| EHMT2 |  |  |
| HNRNPU |  |  |

**Table 3.** Top ten genes according to degree and betweenness centrality in each groups.

| Protein Names | Degree | Protein Names | Betweenness | Common Proteins |
| --- | --- | --- | --- | --- |
| **Normal vs. ACs** |  |  |  |  |
| SRC | 35 | UGCG | 1.00 | SRC |
| ACTA2 | 26 | ABCC4 | 1.00 | ACTA2 |
| HSP90AB1 | 25 | NUDT16L1 | 0.66 | HSP90AB1 |
| EHMT2 | 22 | SRC | 0.22 | CLTC |
| CLTC | 21 | HSP90AB1 | 0.11 | PPIE |
| PPIE | 21 | ACTA2 | 0.10 |  |
| CDKN1A | 21 | CLTC | 0.08 |  |
| RPS6 | 17 | PPIE | 0.07 |  |
| VAMP2 | 17 | HELZ2 | 0.06 |  |
| RBM8A | 17 | STAT1 | 0.06 |  |
| **Normal vs. TSP** |  |  |  |  |
| STAT1 | 18 | STAT1 | 0.27 | STAT1 |
| IFNG | 16 | IFNG | 0.23 | IFNG |
| IRF1 | 14 | CASP3 | 0.17 | IRF1 |
| PSMB8 | 12 | JAK2 | 0.15 | PSMB8 |
| HLA-B | 12 | PSMB8 | 0.07 | HLA-B |
| GBP2 | 11 | IRF1 | 0.07 | JAK2 |
| JAK2 | 9 | HLA-B | 0.05 | GBP2 |
| IFI35 | 9 | PSMB9 | 0.02 |  |
| IFITM1 | 9 | GBP5 | 0.02 |  |
| BST2 | 9 | GBP2 | 0.02 |  |
| **ACs vs. TSP** |  |  |  |  |
| STAT1 | 9 | STAT1 | 0.77 | STAT1 |
| PSME1 | 4 | CASP1 | 0.32 | PSME1 |
| PSMB8 | 4 | PSME1 | 0.17 | PSMB8 |
| CASP1 | 4 | PSMB8 | 0.02 | CASP1 |
| PSME2 | 3 | PSME2 | 0.00 |  |
| IFIT2 | 2 | IFIT2 | 0.00 |  |
| MAPK13 | 2 | MAPK13 | 0.00 |  |
| IL15 | 2 | IL15 | 0.00 |  |
| PTPN6 | 2 | PTPN6 | 0.00 |  |
| AIM2 | 1 | AIM2 | 0.00 |  |

**Figure 1:**
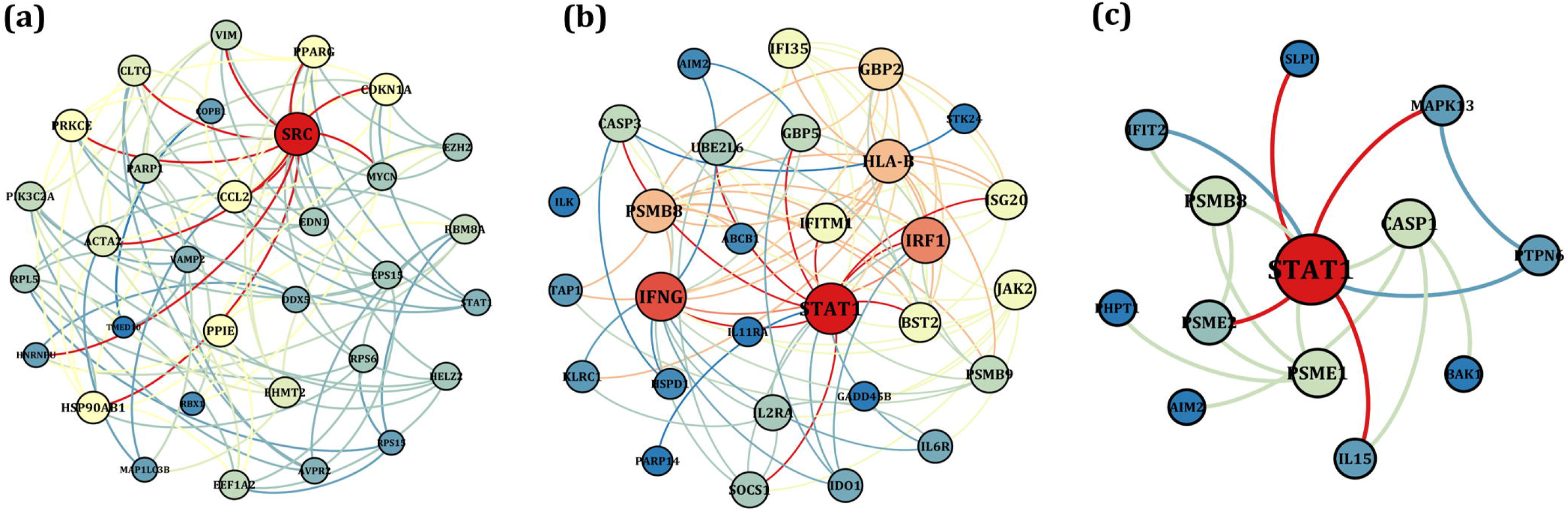
The PPINs constituted between the identified hub DEGs of (a) Normal *vs.* ACs, (b) Normal *vs.* HAM/TSP, (c) ACs *vs.* HAM/TSP groups. The node size and color is indicative of the degree level, so that red color and larger size indicate the higher degree of node.

### Module finding and enriched pathways analysis

To derive the functionality of each sub-networks, the modules were identified by the fast unfolding clustering algorithm. Four, three, and three modules were detected for normal *vs.* ACs, normal *vs.* HAM/TSP, and ACs *vs.* HAM/TSP groups, respectively. The nodes of biologically meaningful modules were enriched in KEGG pathway (Table 4).

**Table 4.** Nodes list of biologically meaningful modules.

| Row | Groups | Genes involved in top module |
| --- | --- | --- |
| 1 | Normal *vs.* ACs | EDN1, SRC, PPARG, PARP1, RBX1, CDKN1A, STAT1, CCL2, VIM, MYCN, EZH2, MAP1LC3B |
| 2 | Normal *vs.* TSP | PSMB8, PSMB9, IRF1, UBE2L6, IFI35, IFITM1, BST2, GBP2, ISG20, HLA-B, TAP1 |
| 3 | ACs *vs.* TSP | PSME1, PSME2, PSMB8, STAT1, IFIT2, PHPT1, SLPI |

In normal *vs.* ACs group, ErbB signaling pathway, Prolactin signaling pathway, MicroRNAs in cancer, Bladder cancer, Transcriptional misregulation in cancer, Chemokine signaling pathway, Hepatitis B, HIF-1 signaling pathway, Pathways in cancer, AGE-RAGE signaling pathway in diabetic complications; in normal *vs.* HAM/TSP group, Viral myocarditis, Autoimmune thyroid disease, Type I diabetes mellitus, Graft-versus-host disease, Primary immunodeficiency, Allograft rejection, Herpes simplex infection, Phagosome, Antigen processing and presentation, and Proteasome; and in ACs *vs.* HAM/TSP group, Thyroid hormone signaling pathway, Toxoplasmosis, Toll-like receptor signaling pathway, Leishmaniasis, AGE-RAGE signaling pathway in diabetic complications, Prolactin signaling pathway, Pancreatic cancer, Inflammatory bowel disease (IBD), Antigen processing and presentation, and Proteasome were enriched (Figure 2).

**Figure 2:**
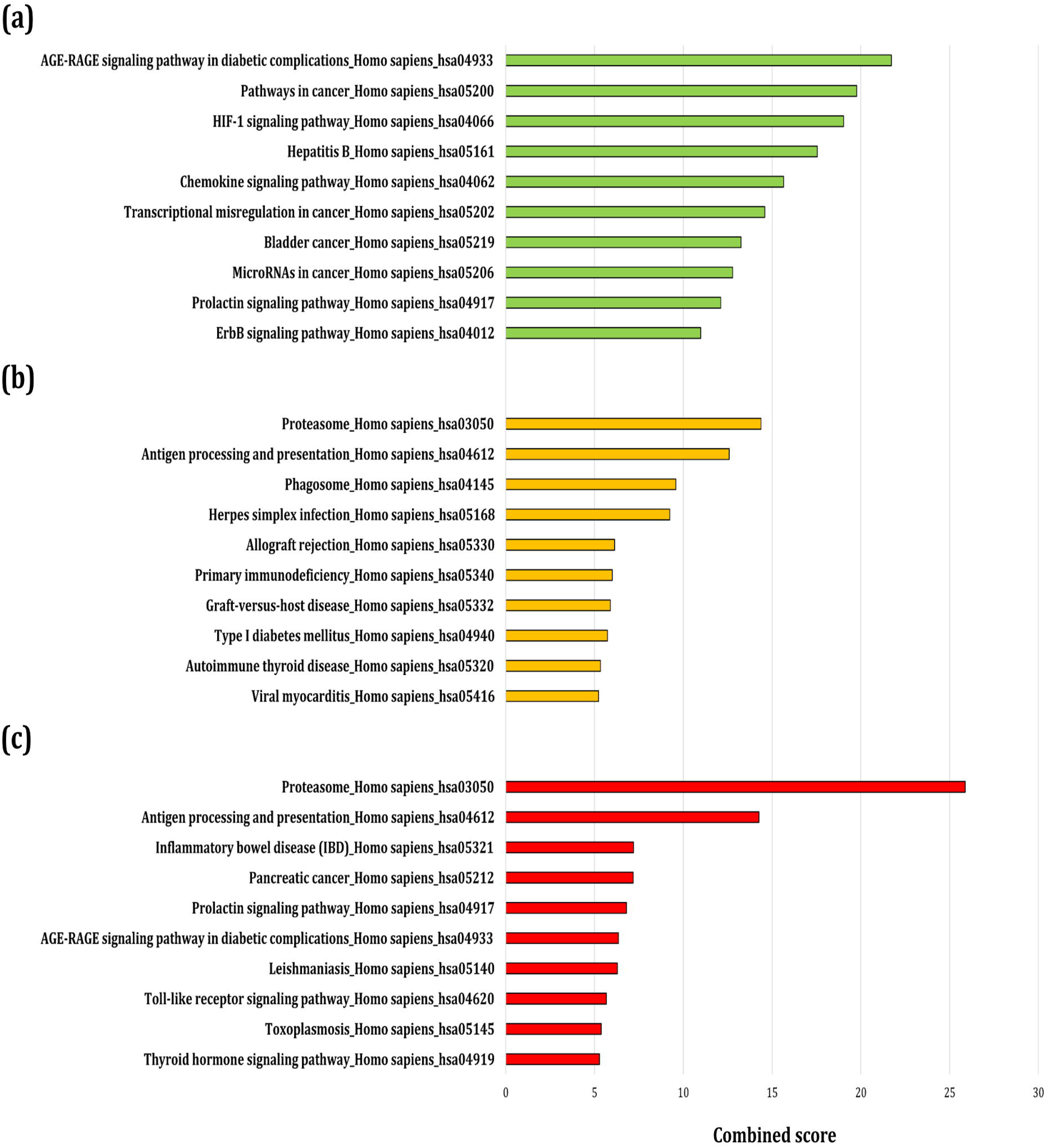
Top ten enriched KEGG pathways of biologically meaningful modules identified from networks of (a) normal *vs.* ACs, (b) Normal *vs.* HAM/TSP, (c) ACs *vs.* HAM/TSP groups.

### Identification of effectual genes in HAM/TSP pathogenesis

To find the most effective genes involved in pathogenesis pathway of HAM/TSP, the meta-analysis results were surveyed in three steps. Firstly, Venn diagram between DEGs of three groups revealed that STAT1 is common between normal *vs.* ACs, normal *vs.* HAM/TSP, and ACs *vs.* HAM/TSP groups; STAT1, GCH1, and TAP1 between normal *vs.* ACs and normal *vs.* HAM/TSP; STAT1, AIM2, PHPT1, PSMB8 between normal *vs.* HAM/TSP, ACs *vs.* HAM/TSP; and STAT1 between normal *vs.* ACs and ACs *vs.* HAM/TSP (Figure 3). Secondly, the super-network of each group was analyzed and top nodes in terms of degree and betweenness were found. The results highlighted SRC, ACTA2, HSP90AB1, CLTC, and PPIE in normal *vs.* ACs group, STAT1, IFNG, IRF1, PSMB8, HLA-B, and JAK2 in normal *vs.* HAM/TSP group, and STAT1, PSME1, PSMB8, CASP1 in ACs *vs.* HAM/TSP group (Table 3). Thirdly, the genes contributed in meaningful modules of sub-networks were determined as EDN1, SRC, PPARG, PARP1, RBX1, CDKN1A, STAT1, CCL2, VIM, MYCN, EZH2, MAP1LC3B in normal *vs.* ACs group, PSMB8, PSMB9, IRF1, UBE2L6, IFI35, IFITM1, BST2, GBP2, ISG20, HLA-B, TAP1 in normal *vs.* HAM/TSP, and PSME1, PSME2, PSMB8, STAT1, IFIT2, PHPT1, and SLPI in ACs *vs.* HAM/TSP network (Table 4). Therefore, STAT1, TAP1, and PSMB8 which were identified in three mentioned analyses were selected for experimental confirmatory tests.

**Figure 3:**
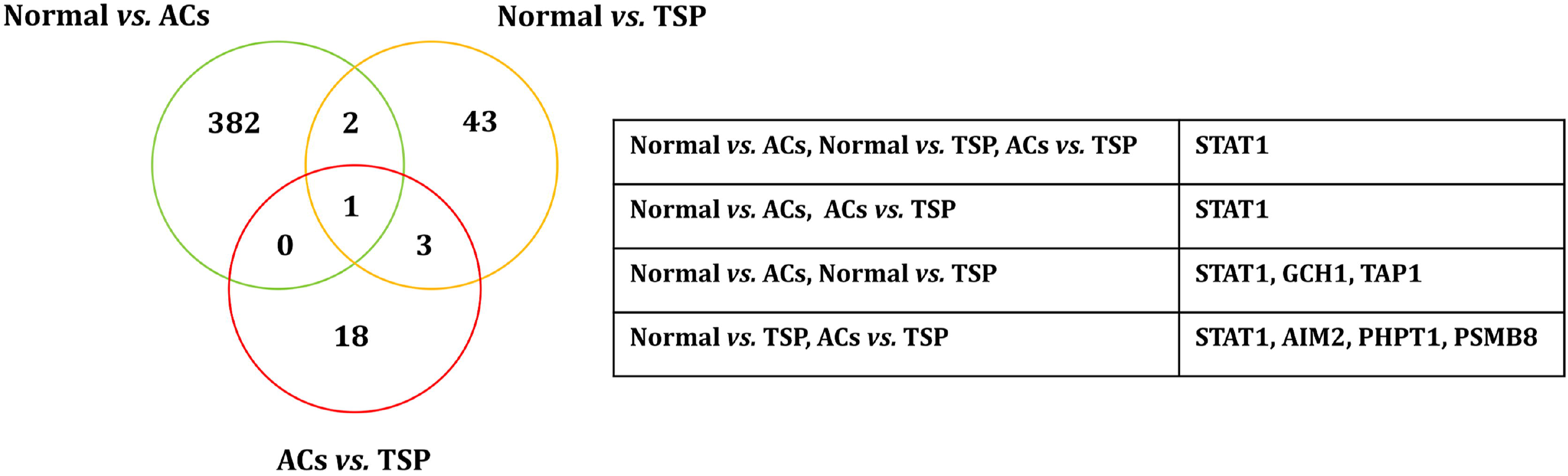
Venn diagram between DEGs of Normal *vs.* ACs, Normal *vs.* HAM/TSP, ACs *vs.* HAM/TSP groups which reveals common genes between groups.

### Demographic data

The mean age of three groups was as follow: normal controls; 41± 2.8, ACs; 42 ± 3.5, HAM/TSP patients; 48 ± 3.6. Any significant difference was found between the ages of three groups.

### Flow Cytometry

Flow Cytometry Data Analyze of T helper and cytotoxic T lymphocytes was done by Flowjo 7.6.1. No significant difference was found among three groups in terms of percentage of T helper (*P*=0.55) and cytotoxic T lymphocytes (*P*=0.12) (Figure 4).

**Figure 4:**
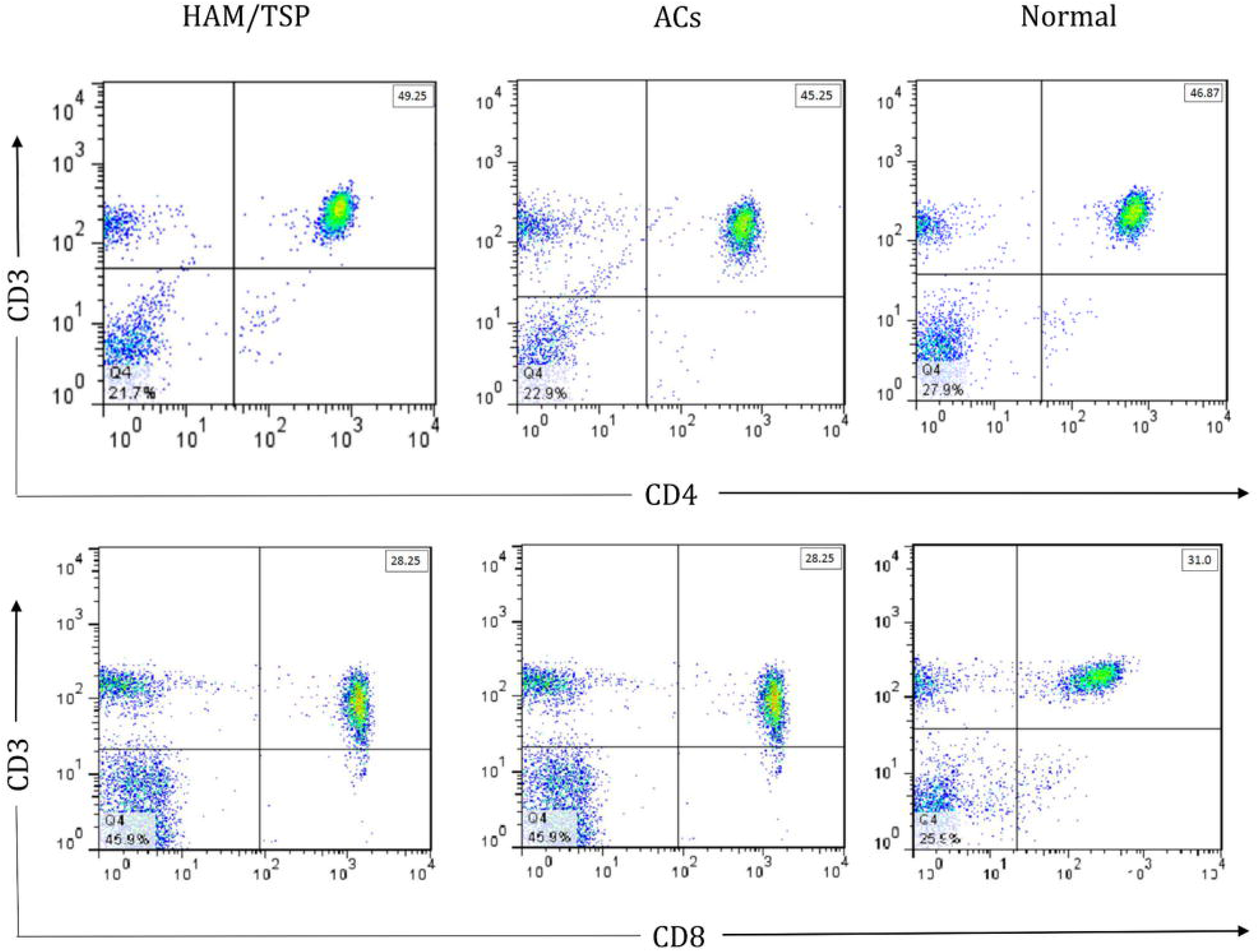
Flow Cytometry Data Analyze of T helper and cytotoxic T lymphocytes.

### HTLV-1 proviral load

All HAM/TSP patients had proviral loads (PVLs) in the range of 216–1160 and all ACs had PVLs in the range of 32–140. The mean PVL of HTLV-1 in the HAM/TSP patients was 455.8 ± 114.7, which was significantly higher (*P*=0.002) than that in the ACs (60.88 ± 12.92) (Figure 5a).

**Figure 5:**
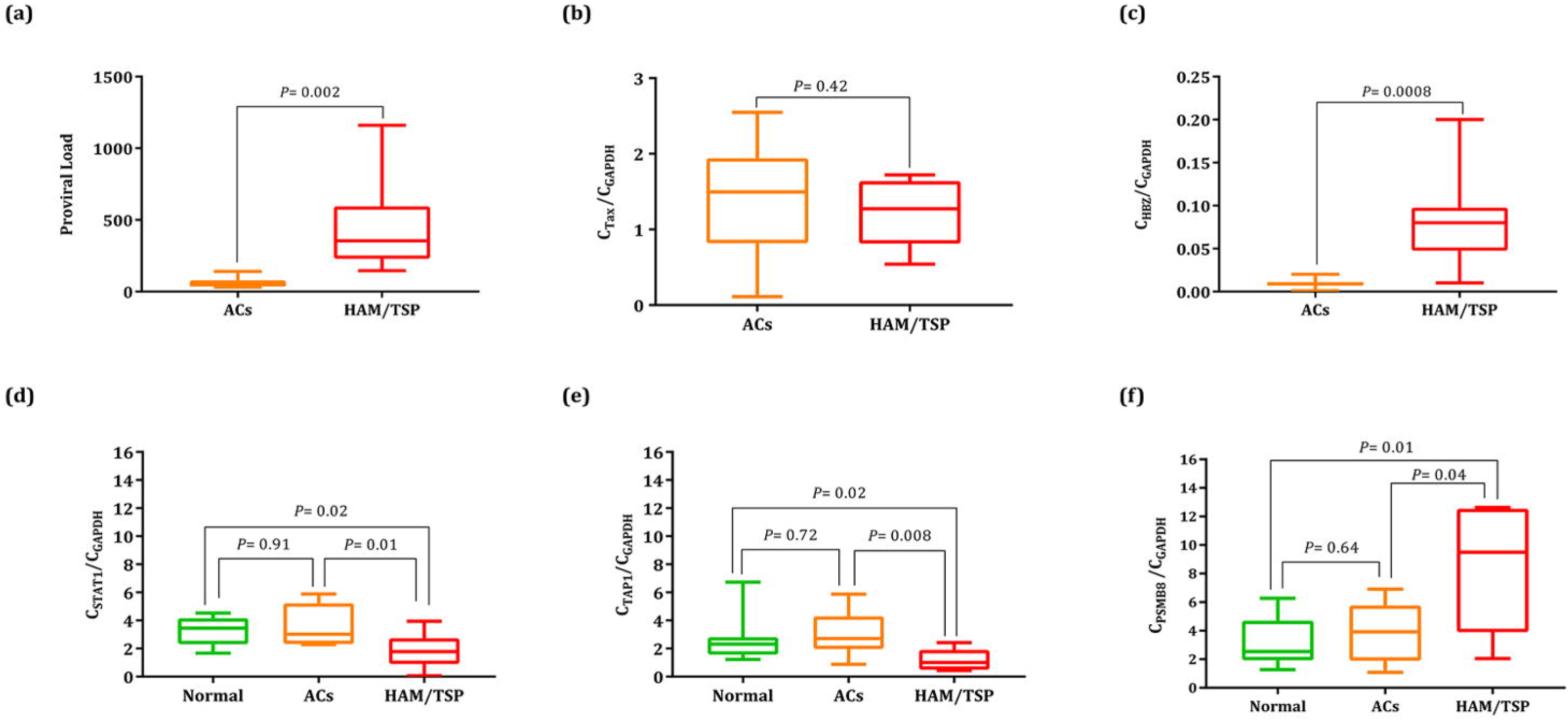
(a) HTLV-I-proviral load. The PVL in HAM/TSP patients was significantly higher than in ACs (*P*□=□0.002). (b) Tax gene expression. No significant difference was found between ACs and HAM/TSP groups (*P*□=□0.42). (c) HBZ gene expressions which was significantly higher in the HAM/TSP group than that in the ACs group (*P*□=□0.0008). (d) STAT1 gene expressions in the Normal, ACs, and HAM/TSP groups. STAT1 gene expression in the HAM/TSP was significantly higher than in Normal (*P*□=□0.02). The STAT1 between AC and HAM/TSP patients was statistically different (*P*□=□0.01). No significant difference was found between Normal and AC patients (*P*□=□0.91). (e) TAP1 gene expressions in the Normal, ACs, and HAM/TSP groups. TAP1 gene expression in the HAM/TSP was significantly higher than in Normal (*P*□=□0.02). The TAP1 between AC and HAM/TSP patients was statistically different (*P*□=□0.008). No significant difference was found between Normal and AC patients (*P*□=□0.72). (e) PSMB8 gene expressions in the Normal, ACs, and HAM/TSP groups. PSMB8 gene expression in the HAM/TSP was significantly higher than in Normal (*P*□=□0.01). The PSMB8 between AC and HAM/TSP patients was statistically different (*P*□=□0.04). No significant difference was found between Normal and AC patients (*P*□=□0.64).

### Real time-quantitative PCR for validation of expression changes

The expression levels of Tax and HBZ were measured in the samples, which revealed the insignificant up-regulation of Tax in ACs group (1.41 ± 0.27) than that in HAM/TSP (1.22 ± 0.16) group (*P*=0.42) and significant higher expression level of HBZ in HAM/TSP group (0.08 ± 0.01) than that in ACs group (0.009 ± 0.001) (*P*=0.0008) (Figure 5b,c). The biomarker potential of STAT1, TAP1, and PSMB8 was surveyed by Real time-quantitative PCR analysis. To this purpose, the differential expressions of genes were analyzed by comparing expression levels in normal, ACs, and HAM/TSP samples. The results revealed the meaningful down-regulation of STAT1 in HAM/TSP (1.8 ± 0.43) samples than those in the AC (3.6 ± 0.52) and normal (3.3 ± 0.36) samples (*P*=0.01 and *P*=0.02, respectively) (Figure 5d). The remarkable down-regulation of TAP1 in HAM/TSP (1.2 ± 0.27) samples than those in the AC (3.0 ± 0.56) and normal (2.7 ± 0.61) samples was observed (*P*=0.008 and *P*=0.02, respectively) (Figure 5e). Also, the expression level of PSMB8 has significantly increased in HAM/TSP (8.5 ± 1.5) samples than those in the AC (3.8 ± 0.74) and normal (3.1 ± 0.61) samples (*P*=0.04 and *P*=0.01, respectively) (Figure 5f). Moreover, the correlation analysis was done to determine the association between different factors. The results indicated the significant correlation between STAT1 and PVL (*P*=0.04, r=0.74) and also between STAT1 and PSMB8 (*P*=0.03, r=0.76) in ACs group. The remarkable associations were observed between Tax and TAP1 (*P*=0.04, r=0.73), STAT1 and PSMB8 (*P*=0.02, r=0.78), HBZ and PVL (*P*=0.05, r=0.70) in HAM/TSP group.

## Discussion

Despite four decades of researches on HTLV-1, many questions remain regarding the pathogenicity mechanism and key proteins involved in various pathological pathways. Moreover, it is also ambiguous that which factors and proteins determine the final destiny of infection by HTLV1 toward HAM/TSP or/and ATLL, while some infected subjects remain in the form of asymptomatic carriers.

Microarray technology is widely used to analyze and measure genomics changes, gene and protein expressions at the high-throughput scale. Despite the high benefits of using this technology, the result of a population cannot be generalized to another population. Data integration and providing a meta-analysis of the reported data improve the validity and reliability of the results. Genomics, transcriptomics, and proteomics data can be combined to obtain more meaningful and appropriate outcomes to find biomarkers and possible pathogenesis pathways (Ramasamy et al., 2008a).

From differential expression analysis of miRNAs samples between normal and ACs groups, four miRNAs including hsa-mir-218, hsa-mir-206, hsa-mir-31, and hsa-mir-34A were identified, which can be considered as biomarkers for diagnosis of AC state.

In complying with previous reports, the identified DEGs implicated in the immune system in HAM/TSP disease. Moreover, the involved molecular network as the primary model was introduced through collection and integration of high-throughput data. The main proteins introduced in this study are STAT1, TAP1, and PSMB8, which were determined based on network analysis, centrality analysis, and module finding.

STAT1 is an important intermediary in responding to IFNs. After binding IFN-I to the cellular receptor, signal transduction occurs through protein kinases which results in the activation of Jak kinase. It, in turn, causes phosphorylation of tyrosine in STAT1 and STAT2. The activated STATs are embedded in the dimer with ISGF3 and IRF9 and enter the nucleus which leads to up-regulation of IFNs and enhances the antiviral response (Zhang et al., 2015;Chen et al., 2017). The significant down-regulation of STAT1 in patients with HAM/TSP was observed compared with asymptomatic carriers and healthy individuals. The decrease in expression of STAT1 is the response of the infected cells to escape HTLV-1 from the immune response associated with HAM/TSP.

The change in expression of STAT1 in ATLL patients has been reported in several studies (Moles et al., 2015). However, no studies have addressed STAT1 expression in HAM/TSP patients. The reduction of STAT1 and subsequent MHC-I in this disease can significantly affect the action of CD8 and NK cells as important cells in the HAM/TSP pathogenesis (Yu et al., 1991;Saito, 2010).

TAP1 is another gene which significantly down-regulated in the HAM/TSP group compared with asymptomatic carriers and normal groups. TAP1 protein which is expressed by the TAP gene involves in the transfer of antigen from the cytoplasm to the endoplasmic reticulum to accompany with MHC-I. HTLV-1 seems to run out from the antiviral response in association with MHC-I due to impairment in the TAP1 function (Bahram et al., 1991). Such occurrence was also observed as a result of infections by other viruses such as EBV, CMV, and adenovirus (Verweij et al., 2015). Similar to STAT1, a decrease in the TAP1 expression can also significantly affect CD8 and NK cells (Yu et al., 1991;Saito, 2010). Therefore, it seems that escaping from CTL-immune response is one of the important mechanism for pathogenicity in HAM/TSP; however, more accurate and detailed studies are needed. In HAM/TSP, the disorder expression of the STAT1 and TAP1 proteins can cause disruption in the immune system.

In addition to two abovementioned genes, a significant increase was observed in the expression of PSMB8 in patients with HAM/TSP in comparison to those who carry the virus and normal subjects. PSMB8 is one of the 17 subunits essential for the synthesis of the 20S proteasome unit (Akiyama et al., 1994). The targeting of proteasome in the HAM/TSP disease is a known mechanism to affect the pathogenicity of HTLV-1 by increasing the activity of genes such as IKBKG, which we have already reported (Mozhgani et al., 2018b). PSMB8 can affect the immune responses because one of the important actions of PSMB8 is involvement in the process of apoptosis (Yang et al., 2009), so its increase in patients with HAM/TSP may be due to this function. Although previous studies reported the role of apoptosis in HAM/TSP pathogenesis (Mozhgani et al., 2018b), there is no comprehensive information regarding the role of PSMB8. In HAM/TSP disease, tissue invasion, cell survival, and inflammation can occur due to the activity of other proteins such as PI3K, PKB, and NFk-B. Also, p53, NF-kappa B, MAPK, and apoptosis pathways are active. PSMB8 and TAP1 in association with PSMB9, IRF1, UBE2L6, IFI35, IFITM1, BST2, GBP2, ISG20, and HLA-B activate the immunodeficiency, autoimmune, and apoptosis pathways. It is noteworthy that the immune mechanisms involved in the HAM/TSP are complicated, so recognition of proteins which have different expressions than the normal group is critical to find the complete pathogenesis pathway (Mozhgani et al., 2018b).

Determining viral factors such as proviral load along with measuring the expression levels of Tax and HBZ genes will be effective in finding the virus action in the patient group. Moreover, host-related factors such as age, family history of the disease, genetics, and host immune status are important (Nagai et al., 1998;Matsuzaki et al., 2001;Okayama et al., 2004;Asquith and Bangham, 2008;Iwanaga et al., 2010;Akbarin et al., 2013).

Destruction of cells in the central nervous system may be due to the release of inflammatory substances from lymphocytes produced by the immune response to contaminated TCD4 + cells, which are referred to as “bystander” damage. It is most likely the mechanism of tissue damage in HAM/TSP disease. In this study, there was no significant difference in the ratio of CD4 to CD8 in the HAM/TSP patients than asymptomatic carriers and healthy subjects; however, a slight increase was observed in the asymptomatic carriers group in comparison to the HAM/TSP and healthy subjects. This may be due to the function of the immunity system to prevent virus replication and progress toward HAM/TSP disease, but more studies with higher sample size are required. Eventually, patients with HAM/TSP have impairment in their immune system induced by the HTLV-1 infection, which includes the innate and adaptive immunity to develop the disease and increase apoptosis (Mozhgani et al., 2018b).

## Conclusion

We employed meta-analysis of high throughput data to find the involved genes in the pathogenesis mechanisms of HAM/TSP disease. Our results confirm that STAT1, TAP1, and PSMB8 are three important genes in HAM/TSP pathogenesis. The up-regulation of PSMB8 in the HAM/TSP subjects enhances the idea that the virus induces apoptosis in the patient’s lymphocytes. Moreover, these proteins in association with other proteins regulate the immune and apoptosis pathways. Finally, the comprehensive studies are needed to increase our knowledge about the pathogenesis pathways and also biomarkers of complex diseases.

## Acknowledgments

Many thanks to the Vice Chancellor for Research, Tehran University of Medical Sciences for supporting the study. This study was the subject of a Ph.D thesis.

## Ethical Approval and Consent to participate

The study was approved by the Ethics Committee of Biomedical Research at TUMS (IR.TUMS.SPH.REC.1396.242).

## Availability of data and material

All relevant data are within the paper.

## Author Contributions Statement

S-HM, MZ-G, MP and MJ performed bioinformatics and statistical analysis. S-HM, MZ-G, MT-R and MJ interpreted and wrote the manuscript. S-HM, NV, TM-A, HF, S-MJ and AK performed experiments. SAR and HR contributed with patient samples. MR performed statistical analysis. MN obtained study funding. MJ, SAR and MN supervised the study. All authors read and approved the final manuscript.

## Conflict of Interest Statement

The authors declare that the research was conducted in the absence of any commercial or financial relationships that could be construed as a potential conflict of interest.

## Contribution to the Field Statement

Human T-lymphotropic virus 1 (HTLV-1) is the first known human retrovirus which is classified as an oncovirus and causes two major human diseases of adult T-cell lymphoma and Human T-lymphotropic virus 1-associated myelopathy/tropical spastic paraparesis (HAM/TSP). Until now, the pathogenicity mechanism and key involved proteins have not been completely identified. Finding main pathogenicity pathway and the involved proteins in the development of disorders help us to disclose the appropriate targets for drug treatments and also vaccine design. Nowadays, by increasing the high-throughput data at genomic, transcriptomic and proteomic levels, it is possible to obtain the phenotype of a particular disease. In this study, we integrated all publically available microarray datasets of HAM/TSP, asymptomatic carriers (ACs), and normal samples to increase the validity of our analysis. The proteins which have special roles in HAM/TSP were identified by statistical analysis. Further network analysis highlighted the different expression of STAT1, TAP1, and PSMB8 which were confirmed by Real-time PCR. Moreover, these proteins in association with other proteins involve in the immune and apoptosis pathways. Therefore, they can be used as a reliable factors for the establishment of pathobiological models and the provision of diagnostic and therapeutic strategies for HAM/TSP.

